# Palmitoylation Couples DLK to JNK3 to Facilitate Pro-degenerative Axon-to-Soma Signaling

**DOI:** 10.1101/2020.11.17.387191

**Authors:** Jingwen Niu, Sabrina M. Holland, Andrea Ketschek, Kaitlin M. Collura, Takashi Hayashi, Gianluca Gallo, Gareth M. Thomas

## Abstract

Dual Leucine-zipper Kinase (DLK, a MAP3K) mediates neuronal responses to diverse injuries and insults via c-Jun N-terminal Kinase (JNK) family Mitogen-activated Protein Kinases (MAPKs). It is unclear why DLK couples to JNKs in mammalian neurons versus other MAPKs, especially because some invertebrate DLK orthologs couple instead to the related p38 family MAPKs. Here we identify two mechanisms that potentially explain this DLK-JNK coupling. First, neural-specific JNK3, but not p38-MAPK, catalyzes positive feedback phosphorylation of DLK that further activates DLK and locks the DLK-JNK3 module in a highly active state. Furthermore, the pro-degenerative JNK2 and JNK3, but not the related JNK1, are endogenously palmitoylated. Moreover, palmitoylation targets both DLK and JNK3 to the same axonal vesicles and JNK3 palmitoylation is essential for pro-degenerative axonal retrograde signaling *in vivo*. These findings provide insights into DLK-JNK signaling relevant to multiple neuropathological conditions and answer long-standing questions regarding the selective pro-degenerative roles of JNK2/3.

## Introduction

Spurred in part by the rising societal burden of age-associated neurodegenerative conditions, there has been intense interest in understanding neuronal signaling mediated by Dual Leucine-zipper Kinase (DLK). DLK is an upstream activator of the response to axonal damage and genetic loss or pharmacological inhibition of DLK strikingly protects neurons against a diverse array of injuries and insults in both the Central and Peripheral Nervous Systems (CNS, PNS) (Fernandes et al. 2014; Le Pichon et al. 2017; Miller et al. 2009; Pozniak et al. 2013; Watkins et al. 2013; Welsbie et al. 2013). However, despite these promising findings, knowledge of DLK-dependent signaling is still incomplete. A better understanding of the molecular control of DLK signaling may help refine current therapeutic strategies and also potentially reveal novel avenues for treatment of diverse neuropathological conditions.

DLK is highly evolutionarily conserved and invertebrate studies have provided important insights regarding DLK roles and regulation (e.g. (Collins et al. 2006; Hammarlund et al. 2009; Nakata et al. 2005; Xiong et al. 2010; Yan et al. 2009)). However, several key questions regarding the control of DLK signaling remain unanswered. For example, it is unclear why DLK (a ‘MAP3K’) couples preferentially to ‘downstream’ c-Jun N-terminal Kinase (JNK) family Mitogen-activated Protein Kinases (MAPKs) in mammalian neurons (Ghosh et al. 2011; Miller et al. 2009), while invertebrate DLK orthologs couple predominantly (in *C. elegans*) or in addition (in *D. melanogaster*) to the related p38 family MAPKs (Klinedinst et al. 2013; Nix et al. 2011; Yan et al. 2009).

A related question regards JNKs themselves – many years ago it was reported that of the three mammalian forms of JNK (JNK1-3), JNK2 and JNK3 (JNK2/3) preferentially mediate neuronal responses to injury and stress, while the related JNK1 instead mediates physiological neuronal regulation (Bjorkblom et al. 2005; Coffey et al. 2002; Yang et al. 1997). These findings have since been broadly replicated in several other models of neuronal injury and degeneration (Brecht et al. 2005; Fernandes et al. 2012; Kuan et al. 2003; Wang et al. 2019). However, the molecular basis for the divergent roles of JNK2/3 versus JNK1 in neurons has not been satisfactorily explained.

We recently reported that covalent modification of DLK with the lipid palmitate, a process called palmitoylation, is critical for somal responses to axonal injury both in cultured neurons and in vivo (Holland et al. 2016; Niu et al. 2020). One provocative finding from this prior work was that palmitoylation not only controls DLK localization but is also critical for DLK to activate JNK3 in non-neuronal cells (Holland et al. 2016), but whether DLK-JNK signaling is similarly palmitoylation-dependent in neurons was not addressed. Another intriguing factor in this regard were prior reports that JNK3 is also palmitoylated (Yang et al. 2012; Yang et al. 2013). Taking these findings together, we hypothesized that palmitoylation of DLK and JNK3 might facilitate preferential coupling of these kinases to drive responses to injury and insult.

In this study we first report that all steps of endogenous DLK-JNK pathway signaling are highly palmitoylation-dependent in neurons. We then report that palmitoylation of DLK is critical to establish a positive feedback loop whereby JNK3 phosphorylates DLK, locking the DLK-JNK3 module in a highly active state. Neither the homologous MAPK p38, nor homologous MAP3Ks, can establish this feedback loop. We further find that JNK2 and JNK3 are both palmitoylated in neurons but the related JNK1 is not. Palmitoylation of JNK3 is not required for JNK3 activation in transfected non-neuronal cells, but is essential to target DLK and JNK3 to the same motile trafficking vesicles in neurons and is critical to traffic JNK3 to optic nerve axons *in vivo*. Finally, using *in vivo* molecular replacement in JNK3 Knockout (KO) mice, we provide evidence that, similar to the importance of DLK palmitoylation (Niu et al. 2020), JNK3 palmitoylation is critical for retrograde signaling following distal optic nerve injury. These findings provide new insights into the control of DLK signaling and answer long-standing questions regarding the molecular basis for selective DLK-JNK coupling and the pro-degenerative roles of JNK2/3.

## Results

### Palmitoylation of DLK is essential for retrograde signaling induced by Trophic Deprivation

We previously reported that palmitoylation of DLK is essential for retrograde signals that result in phosphorylation of the transcription factor c-Jun in neuronal cell bodies after injury to distal axons (Holland et al. 2016). These experiments employed DRG neurons cultured in microfluidic chambers. However, our attempts to gain further insights into how palmitoylation controls DLK-JNK-cJun signals, in particular to determine whether endogenous DLK-JNK kinase activation is palmitoylation-dependent, were initially hampered by the paucity of material available from microfluidic cultures. We therefore took a two-step approach to circumvent this issue. First, we asked whether axon-to-soma signaling in microfluidic cultures in response to Trophic factor Deprivation (TD) similarly requires palmitoylation of DLK. Consistent with this hypothesis, selective withdrawal of Nerve Growth Factor (NGF) from distal axon chambers of microfluidic cultures resulted in robust phosphorylation of c-Jun in DRG neuron cell bodies, which was prevented by DLK shRNA knockdown and rescued by shRNA resistant wild type, but not palmitoyl-mutant, DLK (wtDLK*, DLK-CS*; Fig 1A-C). These findings suggest that DLK palmitoylation is essential for axon-to-soma signaling responses, not just to injury, but to TD and potentially to other axonal insults.

**Figure 1:**
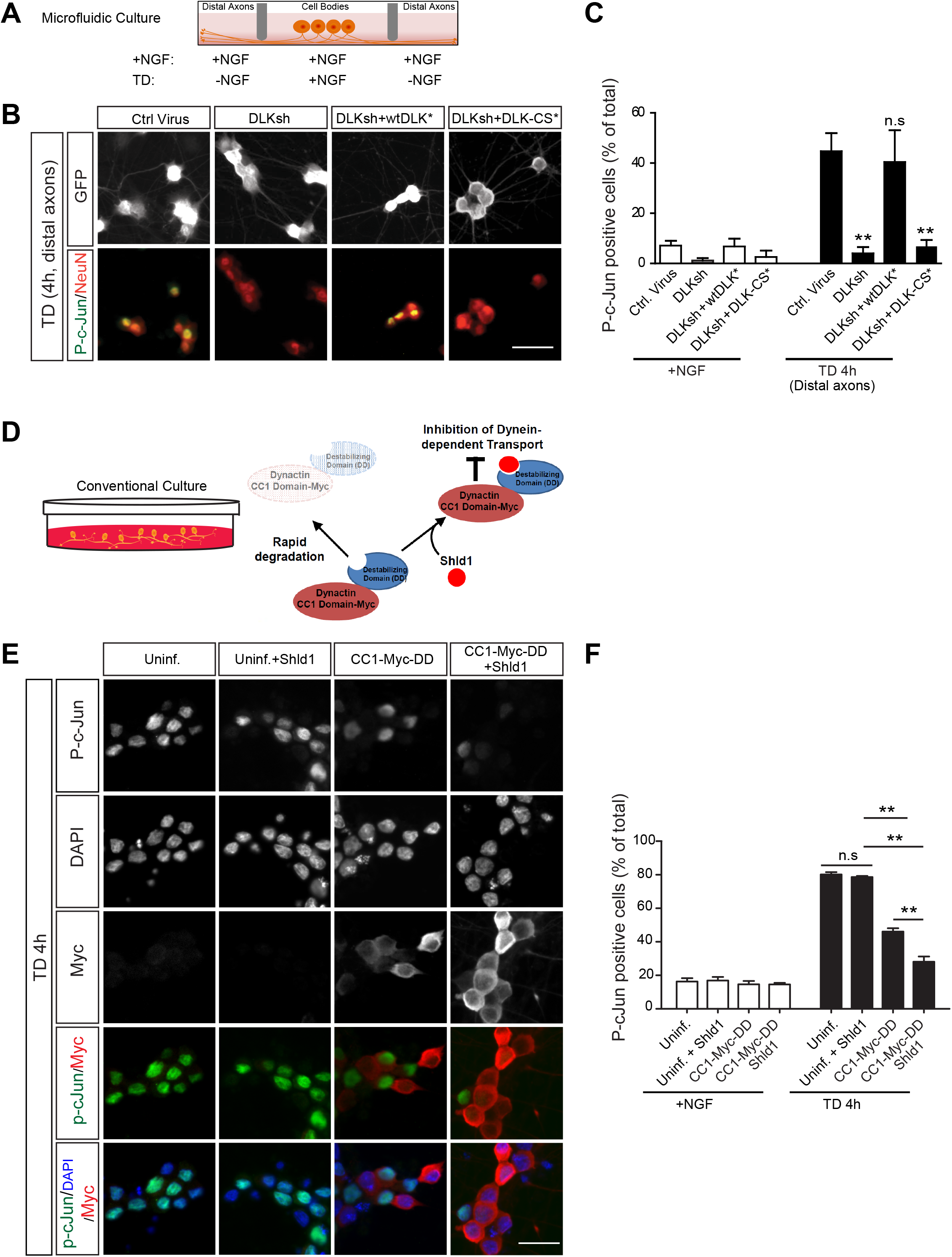
Trophic Deprivation (TD-) induced DLK-JNK-c-Jun signaling in DRG neurons requires palmitoyll DLK and dynein. ***A:*** Schematic of microfluidic culture. Embryonic DRG neurons seeded into central chamber extend axons through microgrooves into distal chambers. Retrograde pro-degenerative signaling can be induced by removing trophic support (Trophic Deprivation, TD) from distal chambers. ***B:*** DRG neurons cultured in microfluidic chambers were infected with the indicated lentiviruses and distal axons were subjected to TD for 4h. Images show cell body/proximal axon chambers immunostained with the indicated antibodies. ***C:*** Quantified data from *B* reveal that TD-induced c-Jun phosphorylation is strongly DLK-dependent and is rescued by shRNA-resistant wtDLK (wtDLK*). Palmitoyl-mutant DLK (DLK-CS*) fails to rescue. **P<0.01 versus control virus TD condition. ANOVA with Tukey *post hoc* test, n=3-4 determinations per condition ***D:*** Schematic of conventional ‘mass’ culture and Destabilization Domain (DD) system. Lentivirally-expressed CC1 domain of dynactin with C-terminal DD and myc tags is normally subject to degradation, keeping expression levels low. Addition of small molecule stabilizer Shld1 increases CC1-myc-DD levels. ***E:*** Mass cultures of DRG neurons were left uninfected or infected with CC1-myc-DD lentivirus and were then treated with Shld1 or vehicle for 24h, prior to global TD for 4h. Cultures were then stained with the indicated antibodies and with DAPI to visualize nuclei. ***F:*** Quantified data from *E* reveal that TD-induced c-Jun phosphorylation is reduced by CC1-myc-DD and further reduced in the presence of Shld1, suggesting that DLK-dependent signaling is dynein-dependent even in conventional culture. ***: p<0.001, n=3 determinations per condition. ANOVA with Bonferroni *post hoc* test.

Second, we asked if TD-induced DLK-JNK signaling in conventionally (‘mass’) cultured DRG neurons requires the retrograde motor protein dynein. If so, this would suggest that similar retrograde signaling mechanisms are likely engaged in both microfluidic and conventional cultures, justifying the use of the latter system (which provides significantly more material) for follow-up biochemical studies. We therefore performed TD in conventionally cultured DRG neurons and in sister cultures lentivirally infected to express the coiled coil 1 (CC1) domain of dynactin, which potently inhibits dynein/dynactin function (Maday et al. 2012). To ensure that we did not disrupt axon development, we used an inducible expression system by fusing the CC1 cDNA to a destabilizing domain (DD; Fig 1D) (Banaszynski et al. 2006), which kept CC1 levels low. We also included a myc tag to detect expression of the CC1-myc-DD fusion protein. The DD domain normally triggers rapid proteasomal degradation of the fusion protein, which can be prevented by adding the otherwise biologically inactive small molecule Shld1 (Fig 1D) (Banaszynski et al. 2006). TD-induced c-Jun phosphorylation was significantly attenuated in cultures expressing CC1-myc-DD, and was further inhibited by adding Shld1 prior to TD (Fig 1D-F). This result suggests that TD-induced DLK signaling is strongly dynein-dependent, even in conventionally cultured DRG neurons.

### All steps of neuronal DLK-JNK-c-Jun signaling are palmitoylation-dependent

Having established the similar involvement of retrograde, palmitoyl-DLK-dependent signals in microfluidic and ‘mass’ cultures, we used the latter system to assess whether endogenous DLK-JNK pathway signaling is palmitoylation-dependent. Consistent with this notion, the palmitoylation inhibitor 2BrP prevented TD-induced phosphorylation of MKK4 (the MAP2K that most likely lies between DLK and JNK in neurons (Itoh et al. 2014)) at sites known to be critical for MKK4 activity (Fig 2A).

**Figure 2:**
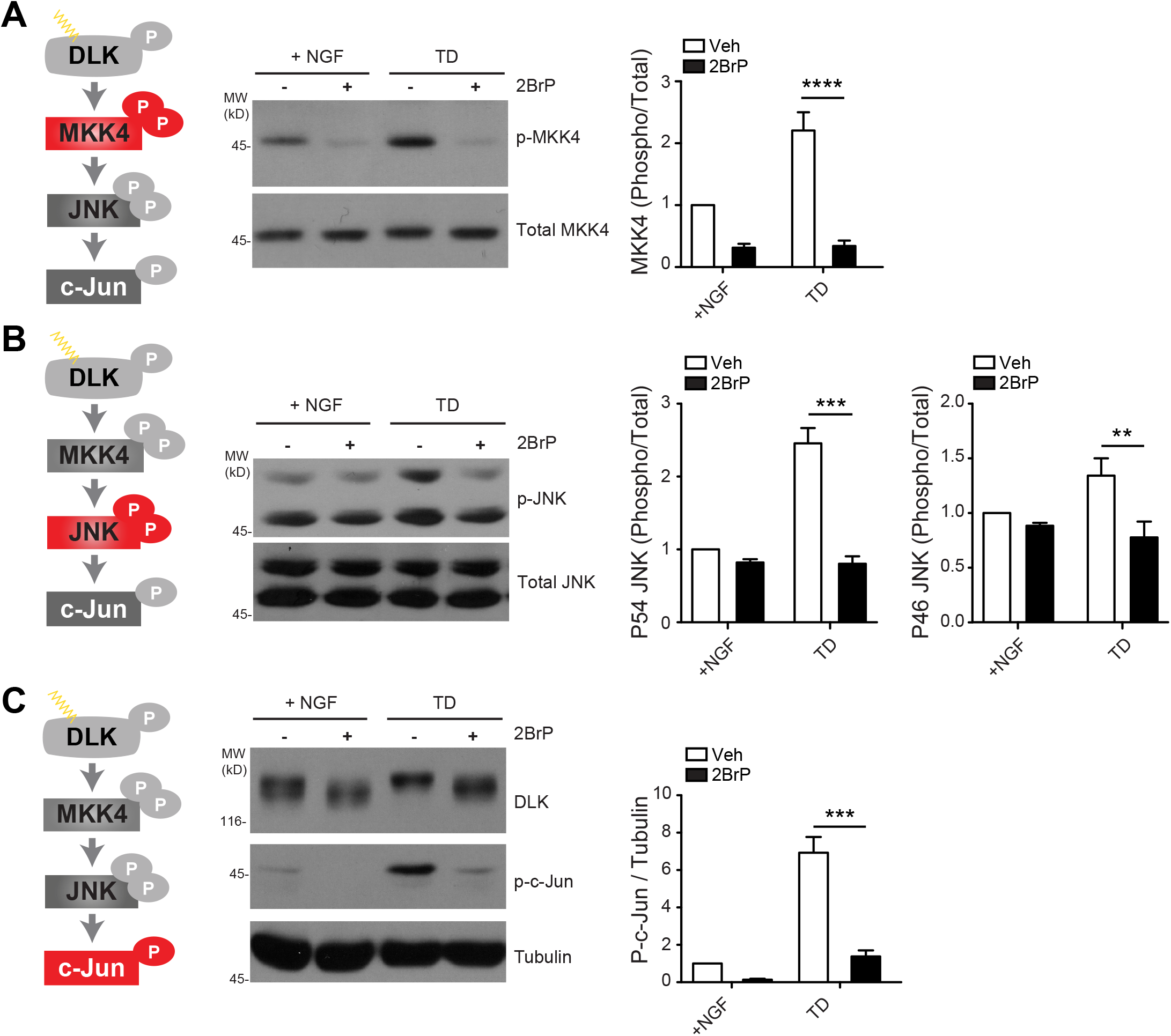
All steps of DLK-JNK Signaling are Palmitoylation-dependent in Trophically Deprived DRG Neurons. ***A:*** DRG neurons were treated with 2BrP (20 μM) or vehicle for 1h and then subjected to TD for 2.5h or left unstimulated in the continued presence of 2BrP or vehicle. Lysates were subjected to SDS-PAGE and subsequent immunoblotting to total and phospho MKK4 (pMKK4). Right panel histogram shows quantified phospho:total MKK4 ratio from n=6 determinations per condition; ****:p<0.0001 versus vehicle treated condition, 2-way ANOVA with Bonferroni post hoc test. ***B:*** As *A*, except that lysates were blotted to detect phospho- and total levels of JNK. Right panel histograms show quantified phospho:total ratios for p54 and p46 forms of JNK, from n=3-6 determinations per condition; ***:p<0.001 versus vehicle treated condition, **:p<0.01 versus vehicle treated condition, 2-way ANOVA with Bonferroni post hoc test. ***C:*** As *A*, expect that lysates were blotted to detect DLK, phospho-c-Jun and tubulin (load control). Right panel shows histogram of phospho-c-Jun:tubulin ratio for n=3-6 determinations per condition; ***:p<0.001 versus vehicle treated condition, 2-way ANOVA with Bonferroni post hoc test. All data are mean ± SEM.

We next asked whether TD-induced activation of JNK is similarly palmitoylation-dependent. There are three JNK genes (JNK1-3, gene names *MAPK8-10*), each of which can undergo alternative splicing events that most notably include or omit a C-terminal extension (Fig S1). The resultant long (p54) and short (p46) forms of JNK1-3 are readily distinguished on SDS-PAGE, but all contain the same activatory Thr-Pro-Tyr motif that is phosphorylated by MKK4 (Barr and Bogoyevitch 2001)(Fig S1). We found that TD-induced phosphorylation of both p46 and p54 JNKs at this Thr-Pro-Tyr motif was palmitoylation-dependent and, interestingly, that the p54 forms of JNK were activated to a greater extent by TD (Fig 2B). TD-induced c-Jun phosphorylation was also highly palmitoylation-dependent (Fig 2C), consistent with our microfluidic experiments and with prior work (Martin et al. 2019). These findings suggest that neuronal DLK-MKK4-JNK signaling is highly palmitoylation-dependent and that effects of TD more potently activate longer (p54) forms of JNK.

In contrast to its blockade of DLK-JNK pathway signaling, 2BrP did not affect either basal phosphorylation or TD-induced dephosphorylation of the pro-survival kinases Akt and ERK (Simon et al. 2016) (Fig S2A, B). Given that 2BrP broadly inhibits cellular palmitoylation and that responses to TD require dynein-dependent transport (Fig 1D, E) we also considered the possibility that effects of 2BrP on DLK-JNK-dependent responses might be due to a generalized impairment in vesicular trafficking. However, 2BrP did not affect either retrograde or anterograde transport of Lysotracker-positive vesicles, which we previously reported co-traffic with DLK ((Holland et al. 2016); Fig S2C). Together, these findings support the interpretation that effects of palmitoylation inhibition on TD-induced c-Jun phosphorylation are due to a direct block of DLK-JNK signaling, rather than to any indirect effects on pro-survival pathways and/or axonal transport.

### Palmitoylation enables feedback phosphorylation and further activation of DLK by JNK3

In the course of our biochemical experiments, we noticed that TD decreased the mobility of endogenous DLK on SDS-PAGE (Fig 1C). A similar TD-induced shift in DLK mobility was reported by (Huntwork-Rodriguez et al. 2013), who suggested that this shift was due to JNK-mediated feedback phosphorylation. Consistent with this interpretation, we observed a similar decrease in DLK mobility when DLK was co-expressed with wild type JNK3 in non-neuronal cells (Fig 3A) and the DLK mobility shift was prevented by treatment of DLK immunoprecipitates with a protein phosphatase (Fig S3). Importantly, genetic or pharmacological block of DLK palmitoylation prevented this phosphorylation-dependent shift in DLK mobility, both in HEK293T cells (Fig 3A, B) and in DRG neurons (Fig 2C). These results suggest that feedback phosphorylation of DLK by JNK is palmitoylation-dependent in both transfected non-neuronal cells and in neurons.

**Figure 3:**
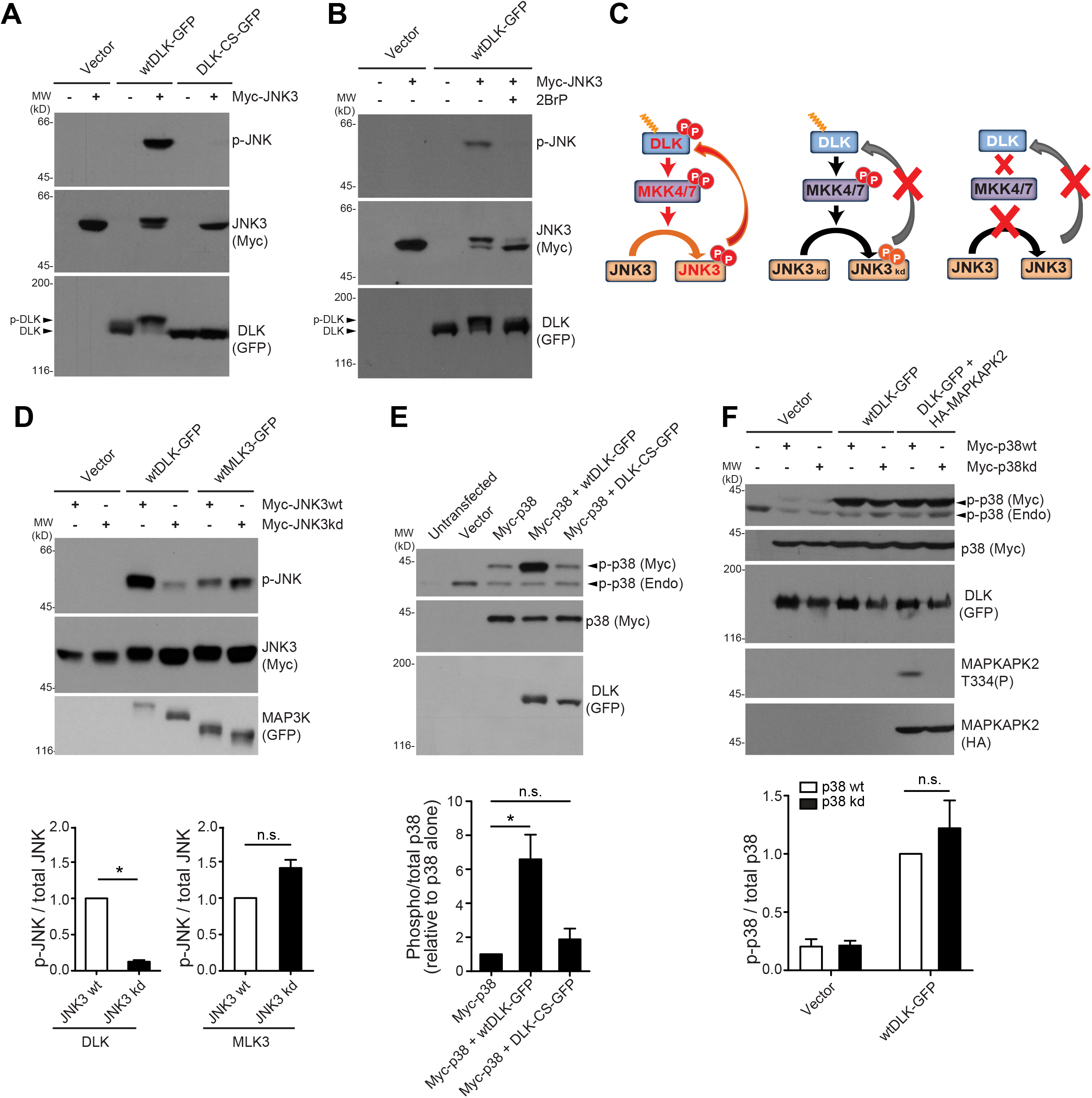
The DLK-JNK3 module engages a palmitoylation-dependent positive feedback loop that further activates DLK. ***A:*** Lysates from HEK293T cells transfected to express the indicated cDNA constructs were blotted to detect the phosphorylated active form of JNK (p-JNK, top), myc (middle) or GFP (bottom). GFP-tagged wild type DLK (wtDLK-GFP), but not palmitoyl mutant DLK (DLK-CS-GFP), greatly increases JNK3 phosphorylation, which correlates with a slowing of DLK-GFP mobility on SDS-PAGE. The slowing of DLK mobility is due to phosphorylation (Compare pDLK and DLK. See also Fig S3). ***B:*** HEK293T cells were transfected with wtDLK-GFP with or without myc-tagged JNK3 (myc-JNK3) and then treated with 2BrP or vehicle. Lysates were blotted as in *A*. Wild type DLK markedly increases JNK3 phosphorylation, which correlates with a marked slowing of wtDLK-GFP mobility on SDS-PAGE (pDLK versus DLK, bottom panel). 2BrP prevents both JNK3 phosphorylation and the change in DLK-GFP mobility. Blots in *A* and *B* are representative of at least three separate experiments. ***C***: Schematic of set-up to determine functional effect of DLK palmitoylation and feedback phosphorylation by JNK3. *Left hand scheme:* Wild type DLK should increase phosphorylation of wild type JNK3, which should in turn phosphorylate DLK. *Middle scheme:* Wild type DLK should be capable of phosphorylating kinase-dead JNK3 (JNK3 kd) but JNK3 kd should be incapable of feedback phosphorylation. *Right hand scheme:* DLK-CS is incapable of driving phosphorylation of JNK3, which in turn cannot phosphorylate DLK. ***D:*** Enabling positive feedback phosphorylation leads to greatly increased phosphorylation of JNK3 by DLK. HEK293T cells were transfected with vector alone, wtDLK-GFP or wtMLK3-GFP (DLK paralog that is not palmitoylated; (Holland et al. 2016)) plus either myc-tagged wild type or kinase dead JNK3 (Myc-JNK3wt, Myc-JNK3kd). Lysates were immunoblotted to detect phosphoJNK (p-JNK, *top*), myc-JNK3 *(middle)* or GFP-tagged MAP3Ks *(bottom)*. JNK3 phosphorylation is markedly higher when feedback loop is engaged (compare top blot, lanes 3, 4). Reduced JNK3 kd phosphorylation is not due to impaired phosphorylatability of the kinase dead JNK3 because phosphorylation of myc-JNK3kd is even more strongly phosphorylated than myc-JNK3wt when MLK3 (MAP3K11) is co-expressed (compare top blot, lanes 5, 6). ***E:*** Quantified data from *D* confirm that phosphorylation of JNK3kd induced by DLK is far lower than that of JNK3wt (*; p=0.0211, n=4, Mann-Whitney test). In contrast, phosphorylation of JNK3kd by MAP3K11 tends toward being greater than that of of JNK3wt (ns; p=0.0636, n=3, Mann-Whitney test). ***F:*** Lack of evidence for positive feedback phosphorylation of DLK by p38 MAPK. HEK293T cells were transfected with myc-tagged wild type or kinase dead p38 MAPK (Myc-p38wt, Myc-p38kd), DLK-GFP and HA-tagged MAPKAPK2 (HA-MAPKAPK2, a downstream p38 MAPK substrate (Rouse et al. 1994)) as indicated. Lysates were blotted to detect phospho–p38 (*top*), myc-p38 (*second row*), DLK-GFP (*third row*), MAPKAPK2 phosphorylated at T334 (p38MAPK phosphorylation site, T334(P), *fourth row*) and HA-tagged MAPKAPK2 (*bottom row*). P38 wildtype does not alter DLK-GFP mobility, compared to p38kd. Cotransfected DLK triggers similar phosphorylation of both p38wt and p38kd. The lower molecular weight band at approximately 38 kDa detected by the p-p38 antibody likely represents endogenous phosphop38 (p-p38 Endo, top panel). MAPKAPK2 T334 signal confirms that myc-p38kd is indeed kinase-dead. **G:** Quantified data from *F* confirm that p38 kinase dead mutation does not affect DLK-induced p38 phosphorylation (ns; p= 0.2817, Mann-Whitney test, n=4). All data are shown as mean ± SEM.

JNK-mediated feedback phosphorylation of DLK correlates with an increase in DLK protein stability following TD (Huntwork-Rodriguez et al. 2013). However, this initial report found that the change in DLK stability only became significant at prolonged times (8h) after TD, and others reported that DLK is stable in DRG neurons over this same 8h timeframe (Karney-Grobe et al. 2018). These findings raised the possibility that JNK-mediated feedback phosphorylation of DLK might instead affect some other aspect of DLK-JNK signaling. Therefore, we considered the possibility that feedback phosphorylation of DLK by JNK might further activate DLK. To test this hypothesis, we compared DLK-dependent phosphorylation of wild type JNK3 (JNK3wt, which can engage the feedback phosphorylation loop) with that of kinase-dead JNK3 (JNK3kd; K55R mutant). In the latter case, JNK3’s activatory Thr-Pro-Tyr motif should still be capable of being phosphorylated by MKK4, but feedback phosphorylation should be prevented, because JNK3kd lacks kinase activity (see schematic, Fig 3C). Interestingly, only co-expressed JNK3wt, but not JNK3kd, decreased DLK mobility on SDS-PAGE (Fig 3D). Moreover, DLK-dependent phosphorylation of JNK3kd at the activatory Thr-Pro-Tyr motif was decreased by >80%, compared to that of JNK3wt (Fig 3D). This result suggests that feedback phosphorylation of DLK by JNK3 further activates DLK, which would thus be predicted to maintain the DLK-JNK3 module in a highly active state.

We next addressed whether similar feedback phosphorylation was observed with other MAP3Ks. The markedly lower phosphorylation of JNK3kd compared to JNK3wt was not seen when MLK3, a non-palmitoylated MAP3K homologous to DLK (Holland et al. 2016), was used as an ‘upstream’ kinase (Fig 3D). This finding suggests that feedback phosphorylation-dependent activation of MAP3K activity is not a general property of this family of kinases. This result also suggests that reduced phosphorylation of JNK3kd by DLK is not because this JNK3 mutant is intrinsically incapable of undergoing phosphorylation, as might be the case if it were, for example, misfolded.

Multiple studies suggest that JNK is the critical downstream MAPK in DLK-dependent signaling in mammalian neurons (Ghosh et al. 2011; Holland et al. 2016; Miller et al. 2009; Simon et al. 2016). However, DLK can also activate the homologous p38 MAPK in cotransfected mammalian cells (Fan et al. 1996) and DLK-dependent effects in *C. elegans* are predominantly mediated by a worm p38 MAPK ortholog (Hammarlund et al. 2009; Yan et al. 2009). Strikingly, although DLK-dependent activation of mammalian p38 MAPK also required DLK palmitoylation (Fig 2E), we observed little, if any, change in DLK mobility induced by p38MAPK co-expression (Fig 3E, F) and no difference between DLK-dependent phosphorylation of wild type and kinase-dead forms of p38 (myc-p38wt, myc-p38kd; Fig 3F). The validity of the p38 MAPK kinase dead mutation was confirmed by the inability of this mutant to phosphorylate its downstream substrate MAPKAP kinase-2 (MAPKAPK2) (Fig 3F; (Rouse et al. 1994)). These findings suggest that while mammalian DLK can activate both p38 and JNK family MAPKs, only the DLK-JNK module engages a positive feedback loop that would be predicted to maintain this module in a highly active state.

### JNK2 and JNK3, but not JNK1, are palmitoylated in DRG neurons

We next asked whether roles of palmitoylation in DLK-JNK signaling are restricted to DLK itself, or might involve control of other elements of the DLK-JNK module. Of the three mammalian JNKs, JNK3 and, to a lesser extent, JNK2, are most implicated in pro-degenerative neuronal signaling (Coffey et al. 2002; Fernandes et al. 2012; Yang et al. 1997). Indeed, JNK3 KO phenocopies loss of DLK itself in multiple neurodegenerative disease models (Fernandes et al. 2012; Fernandes et al. 2014; Pozniak et al. 2013; Yang et al. 1997). We thus asked how DLK might preferentially couple to JNK2/3 compared to other ‘downstream’ MAPKs. Intriguingly, JNK3 itself was reported to be palmitoylated at a C-terminal <u>CC</u>R motif (palmitoylated cysteines underlined) in transfected non-neuronal cells (Yang et al. 2012). We confirmed this finding using a non-radioactive palmitoylation assay, Acyl Biotin Exchange (ABE, Fig 4A) (Thomas et al. 2012; Wan et al. 2007) and also observed robust endogenous JNK3 palmitoylation in DRG neurons (Fig 4B). We then determined the palmitoyl:total levels of the different forms of JNK1-3 in DRG neurons by western blotting ABE fractions and total lysates with specific antibodies (Fig 4C, D, see Fig S4 for antibody validation). Interestingly, approximately 25% of JNK3 and approximately 5% of JNK2’s p54 isoform, which contains a G<u>C</u>R C-terminal motif similar to JNK3’s <u>CC</u>R motif, were palmitoylated (Fig 4C, D). DRG neurons also express p46 isoforms of JNK2 and JNK1 that lack this C-terminal extension containing the palmitoyl-motif (Fig 4C, Fig S1). Consistent with their lack of a palmitoyl-motif, neither p46-JNK1 nor p46-JNK2 was detectable in DRG neuron ABE fractions (Fig 4C). These findings suggest that JNK3 and JNK2, the two forms of JNK most linked to DLK-dependent neuropathology (Fernandes et al. 2012; Fernandes et al. 2014; Pozniak et al. 2013; Yang et al. 1997), are the only palmitoylated JNKs in DRG neurons.

**Figure 4:**
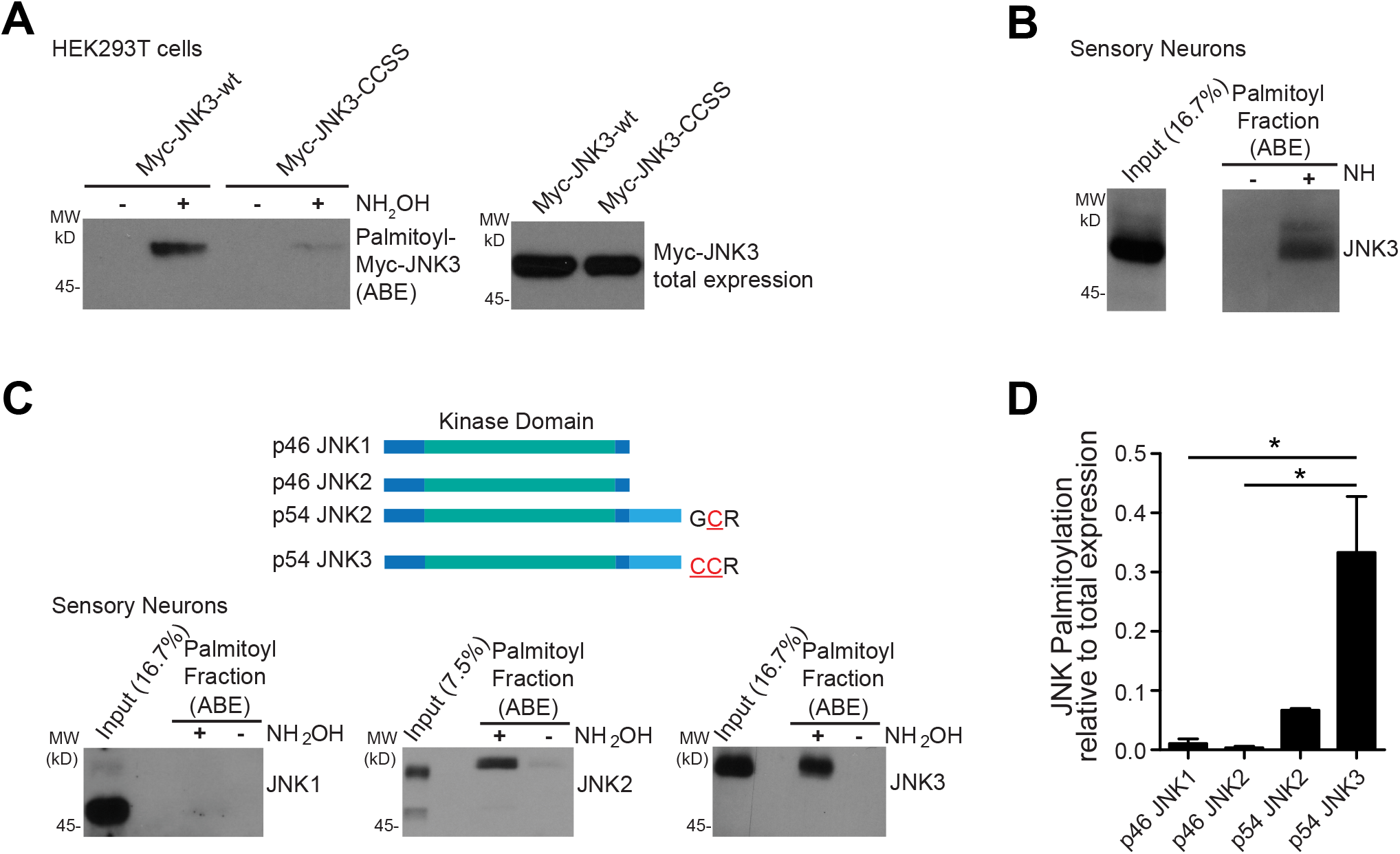
JNK2 and JNK3 are palmitoylated in DRG neurons but JNK1 is not. ***A:*** HEK293T cells were transfected with myc-tagged wild type JNK3 (myc-JNK3-wt) or JNK3 containing a double Cys-Ser mutation at two C-terminal residues (C424S, C425S; ‘myc-JNK3-CCSS’). ABE fractions were blotted to detect palmitoyl-JNK3 and total lysates blotted to detect total JNK3 expression. Myc-JNK3 presence in ABE fractions depends on the key ABE reagent hydroxylamine (NH_2_OH). CCSS mutation greatly reduces JNK3 palmitoylation. Images are representative of three separate experiments. ***B:*** JNK3 is endogenously palmitoylated in DRG DRG neurons. DRG neuron lysates and ABE fractions were blotted to detect JNK3. JNK3 presence in ABE fractions depends on the key ABE reagent hydroxylamine (NH_2_OH). ***C:*** Comparison of JNK1-3 palmitoylation in DRG DRG neurons. The same lysates and ABE fractions from DRG neurons were blotted with specific antibodies to detect JNK1 (left blot) JNK2, (middle blot) and JNK3 (right blot) Antibody specificity is confirmed in Fig S4. Schematic shows forms of JNKs1-3 that are detected in DRG neurons and the likely palmitoyl-motifs of p54-JNK2 and –JNK3 (see also Fig S1). ***D:*** Quantified palmitoyl:total ratios of data from *C* confirm that JNK3 is more highly palmitoylated than other JNKs. N=3-5 determinations per condition. *:p<0.05, one-way ANOVA with Bonferroni *post hoc* test.

### Palmitoylation is dispensable for JNK3 activation in transfected cells but is essential to target JNK3 to DLK-positive vesicles in neurons

We next assessed the potential functional consequence of JNK palmitoylation, focusing on JNK3. In HEK293T cells, JNK3 palmitoyl-site mutation did not significantly affect phosphorylation of JNK3 driven by cotransfected DLK (Fig 5A, B). These findings suggest that palmitoylation is not necessary for JNK3 phosphorylation and activation *per se*.

**Figure 5:**
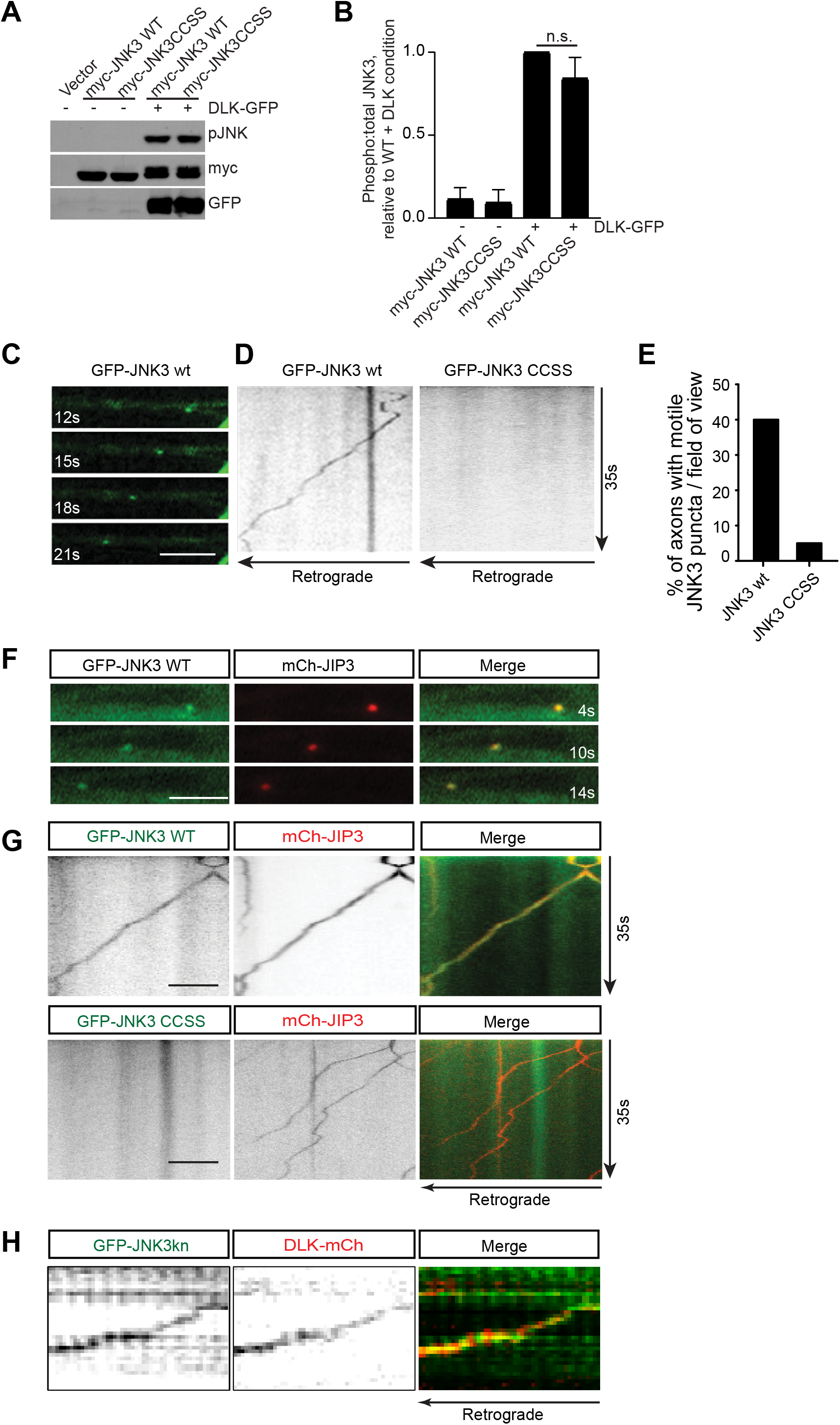
Palmitoylation does not affect JNK3 activation in cotransfected non-neuronal cells but is essential for axonal subcellular localization of JNK3 in cultured neurons. ***A:*** Immunoblots of lysates from HEK293T cells transfected to express the indicated proteins. PhosphoJNK signal, a measure of JNK3 activation, is unaffected by palmitoyl-site (CCSS) mutation. ***B:*** Quantified data from n=3-4 determinations per condition from *A*. n.s.: not significant. ***C:*** Images of axons of DRG neurons lentivirally infected to express GFP-JNK3wt were imaged at the indicated times. GFP-JNK3wt localizes to motile puncta that are likely vesicles (Cavalli et al. 2005). Scale bar: 5 μm. ***D:*** Kymographs generated from DRG axons expressing either GFP-JNK3wt (as in *C*) or GFP-JNK3 CCSS. ***E:*** Quantified data from *D* confirm that JNK3wt is far more frequently detected on motile puncta than JNK3 CCSS (***; p<0.001, Fisher’s exact test). See also Supplementary Movie Files S1, S2. All data are mean ± SEM. ***F, G:*** Individual time-lapse images and kymographs of DRG axons expressing either GFP-JNK3wt or GFP-JNK3 CCSS plus mCh-JIP3. GFP-JNK3wt co-traffics with mCh-JIP3 but GFP-JNK3-CCSS does not. ***H:*** Kymograph of DRG axon expressing GFP-JNK3-KD and DLK-mCh.

The lack of effect of palmitoyl-site mutation on JNK3 activation by cotransfected DLK is also consistent with JNK3’s palmitoyl-motif being distant from its kinase domain, such that it is unlikely to directly impact accessibility by upstream kinases. We therefore asked whether palmitoylation might instead affect JNK3 localization in neurons. Consistent with this notion, GFP-tagged wild type (palmitoylation-competent) JNK3 was detected on motile axonal puncta that are likely vesicles, consistent with a prior report (Fig 5C-E; (Cavalli et al. 2005)). In contrast, a JNK3 palmitoyl-mutant (GFP-JNK3-CCSS) was diffusely distributed and showed no vesicle-like movement (Fig 5D, E). Consistent with their designation as vesicles, axonal puncta of wild type GFP-JNK3 also co-trafficked with an mCherry-tagged version of the vesicle-associated JNK-interacting Protein-3 (mCh-JIP3; Fig 5F, G). In contrast, GFP-JNK3-CCSS remained diffuse in the presence of mCh-JIP3 (Fig 5G). Although palmitoylation was essential for JNK3 vesicular targeting (Fig 5C-G), JNK3 kinase activity was not, because a JNK3 kinase-dead mutant (K55R; ‘GFP-JNK3 KD’) was still detected on motile vesicles (Fig S5). Moreover, GFP-JNK3-KD also co-trafficked with mCherry-tagged DLK (DLK-mCh) on similar axonal puncta, (Fig 4H). These findings suggest that palmitoylation targets both DLK, JNK3 and JIP3 to the same population of axonal vesicles, potentially to facilitate DLK-JNK3 axonal retrograde signaling.

Palmitoylation thus affects JNK3 subcellular localization in cultured DRG neurons, but JNK3-CCSS was still detectable in DRG axons (Fig 5D, G, Movie S2). We reasoned that palmitoylation might even more profoundly affect JNK3 localization *in vivo*, where axonal lengths are considerably longer. To address this question, we first generated Adeno-associated viruses (AAVs) expressing HA-tagged (HA-) JNK3-WT or JNK3-CCSS. We confirmed that AAV-delivered JNK3-WT and JNK3-CCSS expressed at almost identical levels in cultured DRG neurons (Fig S6). However, when we assessed HA-staining after intravitreal delivery of the same HA-JNK3 AAVs to Retinal Ganglion Cells (RGCs), we found that intensity of HA-JNK3-CCSS was significantly greater than that of HA-JNK3-WT in RGC somas but was far less than that of HA-JNK3-WT in the optic nerve (Fig 6A-C). Moreover, in the optic nerve some HA-JNK3-WT was detected as discrete puncta that resembled the axonal vesicles that we detected in cultured neurons (Fig 6C). In contrast, the weak HA-JNK3-CCSS signal in the optic nerve lacked discernible puncta (Fig 6C). These findings suggest that palmitoylation controls JNK3 distribution *in vivo*, likely by targeting JNK3 to axonal vesicles.

**Figure 6:**
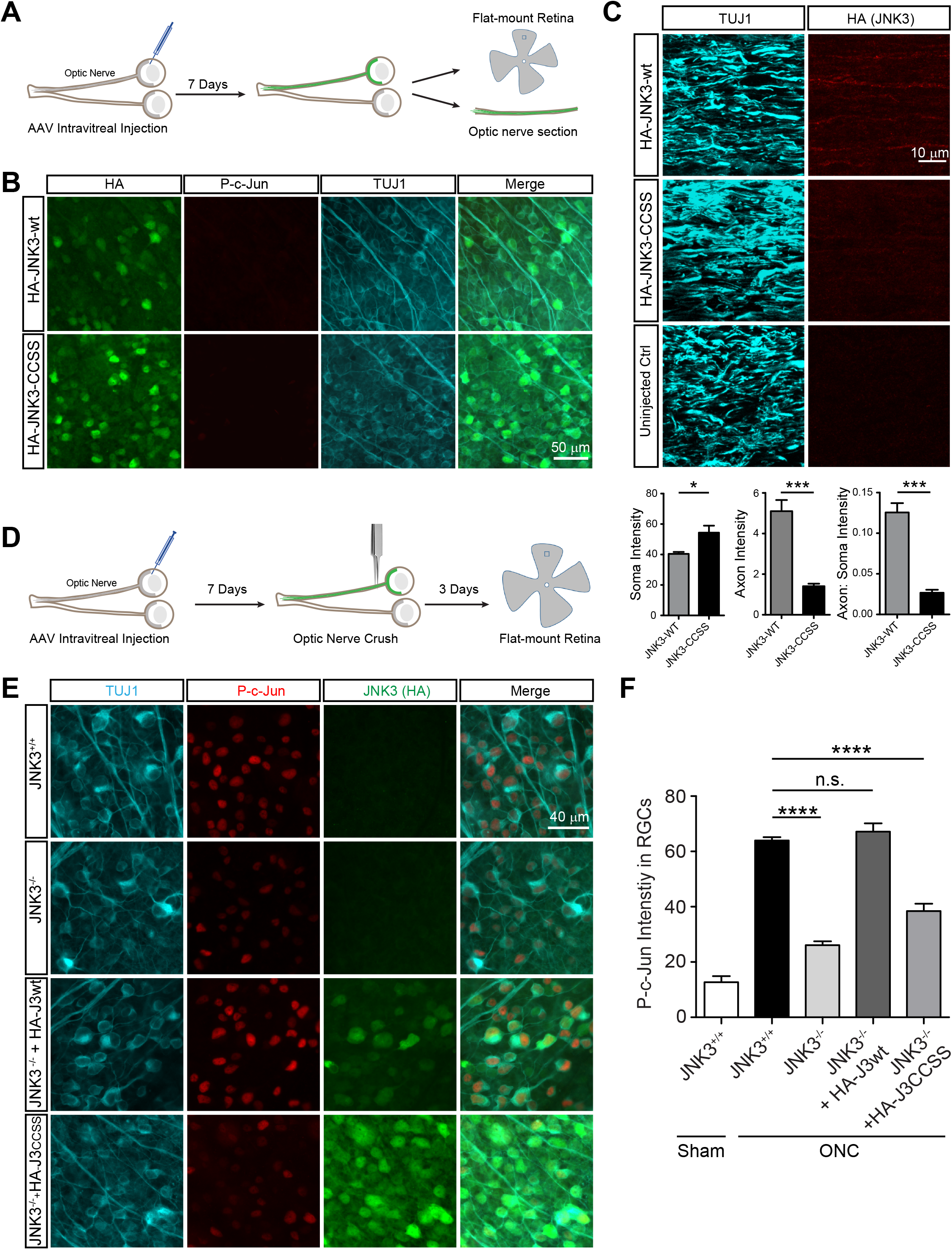
Palmitoylation is critical for JNK3 axonal localization and retrograde injury signaling *in vivo*. ***A:*** Experimental set-up. Blue square indicates approximate region of retina from which images were acquired. ***B:*** Retinal flatmounts from wildtype mice injected with the indicated AAVs and immunostained with the indicated antibodies. ***C:*** Optic nerve sections from the same mice as in *B*, immunostained with the indicated antibodies. Lower panel histograms show quantified somal intensity, optic nerve intensity and optic nerve:somal intensity for the indicated conditions. HA-JNK3wt is much more strongly detected in the optic nerve than HA-JNK3-CCSS. *:p<0.05, ***:p<0.001, t test. ***D:*** Experimental set-up. ***E:*** Retinal flat-mount images from mice of the indicated genotype, injected with the indicated AAVs, subjected to ONC and fixed and immunostained with the indicated antibodies. ***F:*** Quantified data from *E* reveal that ONC-induced c-Jun phosphorylation is reduced in JNK3 KO mice and is rescued by HA-JNK3wt but not HA-JNK3-CCSS. ****:p<0.0001, n.s.: not significant. One-way ANOVA with Bonferroni *post hoc* test, n=3-4 mice per condition.

Finally, we asked whether this differential palmitoylation-dependent distribution might affect retrograde signaling by the DLK-JNK pathway, by assessing the somal response to Optic Nerve Crush (ONC), which requires both palmitoyl-DLK and JNK3 (Fernandes et al. 2012; Fernandes et al. 2014; Niu et al. 2020; Watkins et al. 2013; Welsbie et al. 2013)(Fig 6D). ONC-induced c-Jun phosphorylation was markedly reduced in JNK3 KO mice and was ‘rescued’ by intravitreally injected HA-JNK3-WT AAV, but not by HA-JNK3-CCSS AAVs (Fig 6E, F). These findings suggest that direct palmitoylation is critical for JNK3’s role in axonal retrograde injury signaling *in vivo*.

## Discussion

We previously reported that DLK palmitoylation is critical for initial responses to axonal injury and, unexpectedly, that this modification is also essential for DLK’s kinase activity *in vitro* (Holland et al. 2016; Niu et al. 2020). However, it was unclear from these studies whether palmitoylation controls other aspects of DLK pathway signaling, and why mammalian DLK couples to JNK, rather than to other MAPKs. In this study we identify JNK3 (and likely JNK2) as a key, additional palmitoylated element of the DLK-dependent pro-degenerative signaling pathway and unveil additional aspects of the mechanism and coordination of DLK-JNK signaling.

In particular, JNK2 and JNK3 have been known for many years to play specific roles in neuronal insult and injury signaling (Coffey et al. 2002; Fernandes et al. 2012; Yang et al. 1997), while JNK1 instead mediates physiological regulation (Bjorkblom et al. 2005; Kim et al. 2007). However, no molecular explanation for the selective pathological role of JNK2/3 was previously proposed. More recently JNK2/3 were found to share pro-degenerative roles with DLK (Fernandes et al. 2012; Fernandes et al. 2014), raising the possibility that JNK2 and/or JNK3 act with DLK in a common signaling complex. Our findings reveal that JNK2 and JNK3 are palmitoylated and that palmitoylation of JNK3 is critical for DLK-dependent axonal retrograde signaling (Fig 4, 6). In contrast, JNK1 is not palmitoylated (Fig 4). Palmitoylation thus provides a plausible explanation for the selective pro-degenerative role of JNK2/3.

At the cellular/molecular level, JNK3 was previously reported to localize to and traffic on JIP3-positive axonal vesicles (Cavalli et al. 2005; Drerup and Nechiporuk 2013), but the molecular basis for JNK3’s discrete axonal localization and trafficking also remained unexplained. Our findings that palmitoylation is critical for JNK3 vesicular targeting and for JNK3’s co-trafficking with JIP3 (Fig 5) provide a molecular explanation for these prior findings.

Importantly, palmitoylation is also essential for JNK3 to co-traffic with DLK (Fig 5). This palmitoylation-dependent co-trafficking may be one reason why DLK preferentially couples to JNK, rather than other MAPKs in neurons (Ghosh et al. 2011; Miller et al. 2009). However, an equally important factor may be JNK’s ability to engage a phosphorylation-dependent positive feedback loop that keeps the DLK-JNK module in a highly active state (Fig 3). Neither a homologous MAPK (p38) not a homologous MAP3K (MAP3K11) can engage such a loop, suggesting that this positive feedback may be a unique property of the DLK-JNK module.

Co-trafficking and engagement of a positive feedback loop are thus two key properties of DLK-JNK signaling. Importantly, our additional findings suggest that these two properties may be linked, because although both ‘short’ (p46, lacking C-terminal palmitoylation sites) and ‘long’ (p54, containing palmitoylation sites) forms of JNKs are significantly activated following TD (Fig 2B and (Ghosh et al. 2011)), the effect of TD is more marked for the p54 (palmitoylation-competent) forms (Fig 2B).

Positive feedback phosphorylation and palmitoylation-dependent co-trafficking may also cooperate to convey DLK-JNK3 signals over long distances from distal axons to cell bodies. Importantly, both MKK4 and JNKs are highly sensitive to phosphatase-mediated inactivation (Avdi et al. 2002; Ha et al. 2019; Takekawa et al. 1998), which, if it occurred at any point along their journey, would render the signal conveyed by these kinases meaningless. By maintaining the DLK-JNK3 module in a highly active state, positive feedback phosphorylation of DLK by JNK3 would be predicted to protect the active DLK-JNK3 signal from such phosphatase-mediated inactivation. Such protection may be critical to preserve signal fidelity over the extended times and long distances required for axonal retrograde signaling. The importance of the retrograde motor protein dynein for DLK-JNK signaling (Fig 1) is consistent with this long distance protein co-trafficking model.

Finally, it was surprising that palmitoyl-site mutation affected neither JNK3 activation in transfected non-neuronal cells nor detection of JNK3 in axons of cultured neurons, but was critically important for both JNK3’s axonal localization and axon-to-soma signaling *in vivo*. We recognize that technical explanations could underlie these differences - for example, palmitoyl-site mutation might affect JNK3 activation if JNK3 were expressed at lower levels or for shorter times in non-neuronal cells, or might affect JNK3 axonal targeting in other types of cultured neuron. However, one key point to consider is that certain aspects of palmitoylation-dependent localization and signaling may only be revealed in the intact nervous system, in which the distances involved are often markedly longer than those in culture. Such an explanation is consistent with the critical role of palmitoylation in JNK3 axonal localization and in axon-to-soma signaling in the optic nerve post-injury (Fig 6).

On a practical note, our finding that preventing JNK3 palmitoylation is almost as effective as JNK3 KO in blocking pro-degenerative signaling suggests that this avenue might be pursued therapeutically to reduce the effects of neurodegeneration. The potential of such an approach is increased by findings that signaling by JNK’s upstream activator DLK is also palmitoylation-dependent (Holland et al. 2016; Niu et al. 2020). Our recent finding that the same palmitoyl acyltransferase (PAT), ZDHHC17, regulates not just DLK and likely JNK3, but also the prosurvival factor Nicotinamide Mononucleotide Adenylytransferase-2 (NMNAT2) complicates this issue. However, strategies to selectively prevent DLK-PAT and/or JNK3-PAT interactions, or to enhance DLK/JNK susceptibility to depalmitoylation, nonetheless hold considerable therapeutic promise.

## Materials and Methods

### Animals

Wild type (C57/B6, strain code 000664) and JNK3 KO mice (strain code 004322, on C57/B6 background) were obtained from Jackson labs and were housed in a barrier facility in the Lewis Katz School of Medicine at Temple University. Timed-pregnant female Sprague Dawley rats (Strain code 400, Charles River) used for dissociation of embryonic DRG neurons were sacrificed at E16 as previously described (Holland et al. 2016). All procedures involving animals followed National Institutes of Health guidelines and were approved by the Institutional Animal Care and Use Committee (IACUC) of Temple University.

### AAV Vectors and Preparation

Adeno-associated viruses were made in HEK293T cells by co-transfecting each of the above plasmids with pAAV2 (pACG2)-RC triple mutant (Y444, 500, 730F) (Petrs-Silva et al. 2011) and pHelper (Stratagene, La Jolla, California) plasmids. Cells were lysed 72h post-transfection to release viral particles, which were precipitated using 40% (w/v) polyethylene glycol and purified by cesium chloride density gradient centrifugation. Fractions with refractive index from 1.370 to 1.374 were dialyzed in MWCO 7000 Slide-A-Lyzer cassettes (Thermo Fisher Scientific, Waltham, Massachusetts) overnight at 4°C. AAV titers used for this study were in the range of 1.5-2.5 x 10^12^ genome copies (GC)/ml determined by real-time PCR.

### Lentiviral Vectors and Preparation

Lentiviral vectors carrying DLK shRNA or shRNA-resistant DLK rescue constructs were previously described (Holland 2016). Lentiviral vector FEW (Holland et al. 2016) was modified to accept a sequence (synthesized by Genewiz) coding for a myc epitope tag followed by a Destabilization Domain (DD) based on the F36V L106P mutant of FKBP (Banaszynski et al. 2006). The resulting vector was termed FEW-myc-DD. The CC1 domain of dynactin (generated by PCR from a cDNA template kindly provided by Dr. E. Holzbaur; (Maday et al. 2012)) was subcloned into FEW-myc-DD upstream of the myc epitope, generating FEW-CC1-mycDD. A plasmid coding for mouse JNK3 cDNA (GE Healthcare) was used as a PCR template and the resultant product was subcloned into mammalian expression vector pRK5 downstream of a myc epitope tag, and into a modified version of the lentiviral vector FEW, downstream of a GFP tag. The EF1 alpha promoter in the latter vector was replaced with the human synapsin (hSyn) promoter, generated from Addgene plasmid #26973 by PCR. JNK3 palmitoyl-site mutant (‘CCSS’) was generated by PCR with primers incorporating Cys-Ser mutations. JNK3 kinase dead (K55R mutant) cDNA was generated by Quickchange (Stratagene). mCherry-tagged JIP3 in FEW vector was as previously described (Holland et al. 2016). Mouse p38 (MAPK14) and kinase-dead (K56R) mutant cDNAs were obtained as gBlock fragments (IDT) and subcloned into pRK5 vector downstream of a myc epitope tag. Mouse MAPKAPK2 cDNA was gene synthesized (Genewiz) and sucloned into a modified FEW vector downstream of an N-terminal HA tag. C-terminally GFP-tagged wtDLK and DLK-CS in eGFP-N1 and lentiviral vectors were previously described (Holland et al. 2016).

### Intravitreal Injection, Optic Nerve Crush and Whole mount Retina Immunostaining

Intravitreal injection of AAV and subsequent optic nerve crush were conducted as previously described (Miao et al. 2016). Briefly, mice were anesthetized with 0.01 mg xylazine/g+0.08 mg ketamine/g body weight. 2 μL AAV was injected into the vitreous chamber of 4 week old animals. Optic nerve crush (ONC) was performed 7 days after AAV injection. Mice were transcardially perfused 3 days post-ONC (to detect c-Jun phosphorylation) with 4% PFA/1 x PBS. Eyeballs were subsequently post-fixed for another 2 h prior to dissection of retinas. Whole mount retinal staining was performed as previously described (Miao et al. 2016). Briefly, wide-field images of flat-mounted retinas were acquired using a Nikon 80i epi-fluorescent microscope with a 10x objective.

To quantify c-Jun phosphorylation in AAV-infected retinas, individual RGCs were identified by Tuj1 staining and the nuclear p-c-Jun signal in each RGC was outlined using the signal in the DAPI channel as a mask. Average nuclear p-c-Jun intensity was then quantified using Fiji (Schindelin et al. 2012). For ‘rescue’ experiments using HA-JNK3wt or-SSR mutant, the HA signal in individual RGCs was additionally used to ensure that only AAV-infected RGCs were counted. In each case, 3-4 images were acquired per individual retina and the p-c-Jun intensity in all RGCs per image was averaged to generate a single determination.

### Lentiviral Infection and Trophic Deprivation of Conventional and Microfluidic DRG Neuron cultures

Dorsal root ganglion (DRG) neurons were dissociated from E16 rat embryos and conventional ‘mass’ cultures and microfluidic chamber cultures were prepared as previously described (Holland et al. 2016). DRG neurons in ‘mass’ cultures were infected with lentivirus at two days *in vitro* (DIV4) and subjected to Trophic Deprivation (TD) at DIV10. DRG neurons in microfluidic cultures were infected at DIV1-3 with lentivirus and subjected to TD at DIV 8-10. For all TD experiments, NGF-containing medium was replaced with fresh Neurobasal medium lacking NGF but containing B27 supplement plus 25μg/mL sheep anti-NGF antibody. Neurons were subsequently lysed or fixed at the timepoints indicated in individual Figures.

### Immunofluorescence and imaging in cultured neurons

Dissociated DRG neurons cultured on coverslips were rinsed once with 1x Recording buffer (25mM HEPES pH7.4, 120mM NaCl, 5mM KCl, 2mM CaCl2, 30mM Glucose) and fixed in 4% paraformaldehyde (PFA)/sucrose for 10min at room temperature. Samples were permeabilized in PBS containing 0.25% (w/v) Triton-X-100 for 10 min at 4°C, blocked with PBS containing 10% (v/v) Normal Goat Serum (SouthernBiotech, 0060-01) for 1 hour and incubated in primary antibodies overnight at 4°C in blocking solution. After 3 washes with PBS, cells were incubated for 1 hour at room temperature with AlexaDye-conjugated fluorescent secondary antibodies diluted in blocking solution, prior to 3 final PBS washes and mounting in FluorSave reagent (Millipore Sigma).

c-Jun phosphorylation in cultured DRG neurons was quantified using ImageJ’s cell batch count plugin. Images were thresholded for phospho c-Jun and NeuN signal and the percentage of phospho c-Jun positive cells per total cells (NeuN) was calculated. For experiments involving dynein inhibitor CC1, phospho-c-Jun positive cells per field were counted manually and normalized to the total number of cells per field, assessed with the DNA dye DAPI.

### Live imaging

For all live imaging experiments except Figure 5H, live images of lentivirally infected DRG neurons were acquired at 0.375s intervals using a Zeiss Axio Observer Z1 inverted TIRF microscope with dual wavelength laser system, a-plan-apo TIRF objective (11x, 1.46NA) and Orca R2 camera (Hamamatsu). For images in Fig 5H, lentivirally infected DRG neurons were imaged using a Nikon C2 confocal (60x, 1.3NA) at 2s intervals. Images were analyzed using either Zeiss Axiovision or ImageJ.

### Detection of Palmitoylation by Acyl Biotinyl Exchange (ABE) assay

ABE assays and subsequent western blotting were performed as previously described (Holland et al. 2016).

### CC1 dynein experiments

To determine the dynein dependence of DLK signaling in conventionally cultured DRG neurons, cultures were infected with CC1-mycDD-expressing lentivirus at DIV4 and treated on DIV7 with 1 μM Shld1 (Takara Bio) or vehicle (0.2% (v/v) Methanol) for 24h prior to TD and subsequent fixation and immunostaining as above.

### HEK293T cell feedback phosphorylation experiments

HEK293T cells were transfected using a calcium phosphate based method as described (Holland et al. 2016). 8h post-transfection, cells were washed once with room temperature PBS and then lysed in a solution of 1 volume 5x SDS-PAGE sample buffer (containing fresh 5% (v/v) beta-mercaptoethanol) to 4 volumes of Immunoprecipitation buffer (IPB; (Thomas et al. 2012)) containing fresh Protease Inhibitor Cocktail (Roche) and 1 μM microcystin-LR. Lysates were sonicated, denatured for 5 min at 95°C and subjected to SDS-PAGE and subsequent immunoblotting. In some experiments, 20 μM 2BrP or vehicle control (0.1% v/v ethanol) was added to cells 4h post-transfection.

### Antibodies

Primary antibodies used were as follows: Rabbit anti-phospho c-Jun S63 (#91952, and rabbit anti-phospho c-Jun S63 II (Cell Signaling Technology, #9261, both used for DRG neuron experiments), Rabbit anti-phospho c-Jun Ser-73 (Cell Signaling Technology, #3270, used for RGC experiments); rabbit anti-DLK/MAP3K12 (Sigma/ Prestige, #HPA039936); mouse anti-GFP (Life Technologies, #A11120, clone 3E6 IgG2a), rabbit anti-GFP (Life Technologies, #A11122), mouse anti-Tubulin β3 (BioLegend, TUJ1, Ig2a, Covance catalog# MMS-435P), Myc 9E10 (University of Pennsylvania Cell Center, Catalog #3207), mouse anti-Myc (Enzo, myc9E10), mouse anti-HA11 (BioLegend, 901502, IgG1), rabbit anti HA (Cell Signaling Technology, #3724), mouse anti-tubulin (Millipore Sigma, Catalog #T7451), sheep anti-NGF (CedarLane, catalog #CLMCNET-031), Rabbit anti-phosphoMKK4 (Cell Signaling Technology, #4514, Rabbit anti-MKK4 (Cell Signaling Technology, #9152), Rabbit anti-phosphoAkt T308 (Cell Signaling Technology, #9275, Rabbit anti-Akt (Cell Signaling Technology, #4691), Rabbit anti-phosphoERK (Cell Signaling Technology, #9106), mouse anti-panERK1/2 (Cell Signaling Technology, #9106), mouse anti-phosphoJNK (Cell Signaling Technology, #9255), rabbit anti pan-JNK (Santa Cruz Biotechnology, #SC-571), rabbit anti-phospho-MAPKAPK2 T334 (Cell Signaling Technology, #3041), rabbit anti-phospho-p38 MAPK Cell Signaling Technology, #4511).

### Phosphatase treatment

HEK293T cells were co-transfected with myc-JNK3 plus wtDLK-GFP and lysed in IPB as previously described (Thomas et al. 2012). After centrifugation, soluble supernatants were immunopreciptated with monoclonal anti-GFP antibody precoupled to Protein G Sepharose beads and immunoprecipitates were washed extensively with IPB containing 0.5M NaCl followed by further washes with IPB and then with lambda phosphatase assay buffer (NEB). Equal volumes of each immunoprecipitated sample were then aliquoted into two tubes and beads were then resuspended with 100 microliters of lambda phosphatase assay buffer containing 400 units of lambda protein phosphatase (NEB), or with 100 microliters of lambda phosphatase assay buffer alone. Samples were incubated for 30 min at 30°C with regular flick-mixing prior to addition of one-fifth volume of 5x SDS-PAGE sample buffer. Samples were then denatured for 5 min at 95°C and subjected to SDS-PAGE and subsequent western blotting.

## Supporting information

JNK3 MS Suppl Figs n Legends

## Acknowledgments

We thank Dr. E Holzbaur (UPenn) for CC1 cDNA, Dr. George Smith for overseeing AAV virus preparation, Drs. Heykyeong Jeong and Shaun Sanders for advice on optic nerve sectioning and statistical analysis, respectively, and Dr. C. Su (UC San Diego) for invaluable discussions. This work was supported by grants from National Institutes of Health (R01NS094402) and Shriners Hospitals for Children Grants (85600 PHI) to G.M.T. AAV production was made possible by a Special Shared Facilities grant for viral production from Shriners Hospitals fro Children.

## Author Contributions

Conceptualization, Supervision and Funding Acquisition: G.M.S., G.G., G.M.T.

Investigation: J.N., S.M.H., A.K., K.M.C., T.H., G.M.T.

Methodology: J.N., S.M.H., G.M.T.

Resources: G.M.S., G.G., G.M.T.

## References

Avdi NJ, Malcolm KC, Nick JA, Worthen GS. 2002. A role for protein phosphatase-2A in p38 mitogen-activated protein kinase-mediated regulation of the c-Jun NH(2)-terminal kinase pathway in human neutrophils. J Biol Chem 277(43):40687–40696.

Banaszynski LA, Chen LC, Maynard-Smith LA, Ooi AG, Wandless TJ. 2006. A rapid, reversible, and tunable method to regulate protein function in living cells using synthetic small molecules. Cell 126(5):995–1004.

Barr RK, Bogoyevitch MA. 2001. The c-Jun N-terminal protein kinase family of mitogen-activated protein kinases (JNK MAPKs). Int J Biochem Cell Biol 33(11):1047–1063.

Bjorkblom B, Ostman N, Hongisto V, Komarovski V, Filen JJ, Nyman TA, Kallunki T, Courtney MJ, Coffey ET. 2005. Constitutively active cytoplasmic c-Jun N-terminal kinase 1 is a dominant regulator of dendritic architecture: role of microtubule-associated protein 2 as an effector. J Neurosci 25(27):6350–6361.

Brecht S, Kirchhof R, Chromik A, Willesen M, Nicolaus T, Raivich G, Wessig J, Waetzig V, Goetz M, Claussen M, Pearse D, Kuan CY, Vaudano E, Behrens A, Wagner E, Flavell RA, Davis RJ, Herdegen T. 2005. Specific pathophysiological functions of JNK isoforms in the brain. Eur J Neurosci 21(2):363–377.

Cavalli V, Kujala P, Klumperman J, Goldstein LS. 2005. Sunday Driver links axonal transport to damage signaling. J Cell Biol 168(5):775–787.

Coffey ET, Smiciene G, Hongisto V, Cao J, Brecht S, Herdegen T, Courtney MJ. 2002. c-Jun N-terminal protein kinase (JNK) 2/3 is specifically activated by stress, mediating c-Jun activation, in the presence of constitutive JNK1 activity in cerebellar neurons. J Neurosci 22(11):4335–4345.

Collins CA, Wairkar YP, Johnson SL, DiAntonio A. 2006. Highwire restrains synaptic growth by attenuating a MAP kinase signal. Neuron 51(1):57–69.

Drerup CM, Nechiporuk AV. 2013. JNK-interacting protein 3 mediates the retrograde transport of activated c-Jun N-terminal kinase and lysosomes. PLoS Genet 9(2):e1003303.

Fan G, Merritt SE, Kortenjann M, Shaw PE, Holzman LB. 1996. Dual leucine zipper-bearing kinase (DLK) activates p46SAPK and p38mapk but not ERK2. J Biol Chem 271(40):24788–24793.

Fernandes KA, Harder JM, Fornarola LB, Freeman RS, Clark AF, Pang IH, John SW, Libby RT. 2012. JNK2 and JNK3 are major regulators of axonal injury-induced retinal ganglion cell death. Neurobiol Dis 46(2):393–401.

Fernandes KA, Harder JM, John SW, Shrager P, Libby RT. 2014. DLK-dependent signaling is important for somal but not axonal degeneration of retinal ganglion cells following axonal injury. Neurobiol Dis 69:108–116.

Ghosh AS, Wang B, Pozniak CD, Chen M, Watts RJ, Lewcock JW. 2011. DLK induces developmental neuronal degeneration via selective regulation of proapoptotic JNK activity. J Cell Biol 194(5):751–764.

Ha J, Kang E, Seo J, Cho S. 2019. Phosphorylation Dynamics of JNK Signaling: Effects of DualSpecificity Phosphatases (DUSPs) on the JNK Pathway. Int J Mol Sci 20(24).

Hammarlund M, Nix P, Hauth L, Jorgensen EM, Bastiani M. 2009. Axon regeneration requires a conserved MAP kinase pathway. Science 323(5915):802–806.

Holland SM, Collura KM, Ketschek A, Noma K, Ferguson TA, Jin Y, Gallo G, Thomas GM. 2016. Palmitoylation controls DLK localization, interactions and activity to ensure effective axonal injury signaling. Proc Natl Acad Sci U S A 113(3):763–768.

Huntwork-Rodriguez S, Wang B, Watkins T, Ghosh AS, Pozniak CD, Bustos D, Newton K, Kirkpatrick DS, Lewcock JW. 2013. JNK-mediated phosphorylation of DLK suppresses its ubiquitination to promote neuronal apoptosis. J Cell Biol 202(5):747–763.

Itoh T, Horiuchi M, Ikeda RH, Jr., Xu J, Bannerman P, Pleasure D, Penninger JM, Tournier C, Itoh A. 2014. ZPK/DLK and MKK4 Form the Critical Gateway to Axotomy-Induced Motoneuron Death in Neonates. J Neurosci 34(32):10729–10742.

Karney-Grobe S, Russo A, Frey E, Milbrandt J, DiAntonio A. 2018. HSP90 is a chaperone for DLK and is required for axon injury signaling. Proc Natl Acad Sci U S A.

Kim MJ, Futai K, Jo J, Hayashi Y, Cho K, Sheng M. 2007. Synaptic accumulation of PSD-95 and synaptic function regulated by phosphorylation of serine-295 of PSD-95. Neuron 56(3):488–502.

Klinedinst S, Wang X, Xiong X, Haenfler JM, Collins CA. 2013. Independent pathways downstream of the Wnd/DLK MAPKKK regulate synaptic structure, axonal transport, and injury signaling. J Neurosci 33(31):12764–12778.

Kuan CY, Whitmarsh AJ, Yang DD, Liao G, Schloemer AJ, Dong C, Bao J, Banasiak KJ, Haddad GG, Flavell RA, Davis RJ, Rakic P. 2003. A critical role of neural-specific JNK3 for ischemic apoptosis. Proc Natl Acad Sci U S A 100(25):15184–15189.

Maday S, Wallace KE, Holzbaur EL. 2012. Autophagosomes initiate distally and mature during transport toward the cell soma in primary neurons. J Cell Biol 196(4):407–417.

Martin DDO, Kanuparthi PS, Holland SM, Sanders SS, Jeong HK, Einarson MB, Jacobson MA, Thomas GM. 2019. Identification of Novel Inhibitors of DLK Palmitoylation and Signaling by High Content Screening. Sci Rep 9(1):3632.

Miao L, Yang L, Huang H, Liang F, Ling C, Hu Y. 2016. mTORC1 is necessary but mTORC2 and GSK3beta are inhibitory for AKT3-induced axon regeneration in the central nervous system. Elife 5:e14908.

Miller BR, Press C, Daniels RW, Sasaki Y, Milbrandt J, DiAntonio A. 2009. A dual leucine kinase-dependent axon self-destruction program promotes Wallerian degeneration. Nat Neurosci 12(4):387–389.

Nakata K, Abrams B, Grill B, Goncharov A, Huang X, Chisholm AD, Jin Y. 2005. Regulation of a DLK-1 and p38 MAP kinase pathway by the ubiquitin ligase RPM-1 is required for presynaptic development. Cell 120(3):407–420.

Niu J, Sanders SS, Jeong HJ, Holland SM, Sun Y, Collura KM, Hernandez LM, Huang H, Hayden MR, Smith GM, Hu Y, Jin Y, Thomas GM. 2020. Coupled control of distal axon integrity and somal responses to axonal damage by the palmitoyl acyltransferase ZDHHC17. Cell Reports, 33(7) in press. doi: 10.1016/j.celrep.2020.108365

Nix P, Hisamoto N, Matsumoto K, Bastiani M. 2011. Axon regeneration requires coordinate activation of p38 and JNK MAPK pathways. Proc Natl Acad Sci U S A 108(26):10738–10743.

Petrs-Silva H, Dinculescu A, Li Q, Deng WT, Pang JJ, Min SH, Chiodo V, Neeley AW, Govindasamy L, Bennett A, Agbandje-McKenna M, Zhong L, Li B, Jayandharan GR, Srivastava A, Lewin AS, Hauswirth WW. 2011. Novel properties of tyrosine-mutant AAV2 vectors in the mouse retina. Mol Ther 19(2):293–301.

Pozniak CD, Sengupta Ghosh A, Gogineni A, Hanson JE, Lee SH, Larson JL, Solanoy H, Bustos D, Li H, Ngu H, Jubb AM, Ayalon G, Wu J, Scearce-Levie K, Zhou Q, Weimer RM, Kirkpatrick DS, Lewcock JW. 2013. Dual leucine zipper kinase is required for excitotoxicity-induced neuronal degeneration. J Exp Med 210(12):2553–2567.

Rouse J, Cohen P, Trigon S, Morange M, Alonso-Llamazares A, Zamanillo D, Hunt T, Nebreda AR. 1994. A novel kinase cascade triggered by stress and heat shock that stimulates MAPKAP kinase-2 and phosphorylation of the small heat shock proteins. Cell 78(6):1027–1037.

Schindelin J, Arganda-Carreras I, Frise E, Kaynig V, Longair M, Pietzsch T, Preibisch S, Rueden C, Saalfeld S, Schmid B, Tinevez JY, White DJ, Hartenstein V, Eliceiri K, Tomancak P, Cardona A. 2012. Fiji: an open-source platform for biological-image analysis. Nat Methods 9(7):676–682.

Simon DJ, Pitts J, Hertz NT, Yang J, Yamagishi Y, Olsen O, Mark MT, Molina H, Tessier-Lavigne M. 2016. Axon Degeneration Gated by Retrograde Activation of Somatic Pro-apoptotic Signaling. Cell 164(5):1031–1045.

Takekawa M, Maeda T, Saito H. 1998. Protein phosphatase 2Calpha inhibits the human stress-responsive p38 and JNK MAPK pathways. Embo J 17(16):4744–4752.

Thomas GM, Hayashi T, Chiu SL, Chen CM, Huganir RL. 2012. Palmitoylation by DHHC5/8 targets GRIP1 to dendritic endosomes to regulate AMPA-R trafficking. Neuron 73(3):482–496.

Wan J, Roth AF, Bailey AO, Davis NG. 2007. Palmitoylated proteins: purification and identification. Nat Protoc 2(7):1573–1584.

Wang RR, Li CF, Wang DZ, Zhang CW, Liu GX. 2019. c-Jun N-terminal kinase 3 deficiency protects axotomized retinal ganglion cells via affecting mitochondria involved apoptosis pathway. Int J Ophthalmol 12(1):30–37.

Watkins TA, Wang B, Huntwork-Rodriguez S, Yang J, Jiang Z, Eastham-Anderson J, Modrusan Z, Kaminker JS, Tessier-Lavigne M, Lewcock JW. 2013. DLK initiates a transcriptional program that couples apoptotic and regenerative responses to axonal injury. Proc Natl Acad Sci U S A 110(10):4039–4044.

Welsbie DS, Yang Z, Ge Y, Mitchell KL, Zhou X, Martin SE, Berlinicke CA, Hackler L, Jr., Fuller J, Fu J, Cao LH, Han B, Auld D, Xue T, Hirai S, Germain L, Simard-Bisson C, Blouin R, Nguyen JV, Davis CH, Enke RA, Boye SL, Merbs SL, Marsh-Armstrong N, Hauswirth WW, DiAntonio A, Nickells RW, Inglese J, Hanes J, Yau KW, Quigley HA, Zack DJ. 2013. Functional genomic screening identifies dual leucine zipper kinase as a key mediator of retinal ganglion cell death. Proc Natl Acad Sci U S A 110(10):4045–4050.

Xiong X, Wang X, Ewanek R, Bhat P, Diantonio A, Collins CA. 2010. Protein turnover of the Wallenda/DLK kinase regulates a retrograde response to axonal injury. J Cell Biol 191(1):211–223.

Yan D, Wu Z, Chisholm AD, Jin Y. 2009. The DLK-1 kinase promotes mRNA stability and local translation in C. elegans synapses and axon regeneration. Cell 138(5):1005–1018.

Yang DD, Kuan CY, Whitmarsh AJ, Rincon M, Zheng TS, Davis RJ, Rakic P, Flavell RA. 1997. Absence of excitotoxicity-induced apoptosis in the hippocampus of mice lacking the Jnk3 gene. Nature 389(6653):865–870.

Yang G, Liu Y, Yang K, Liu R, Zhu S, Coquinco A, Wen W, Kojic L, Jia W, Cynader M. 2012. Isoform-specific palmitoylation of JNK regulates axonal development. Cell Death Differ 19(4):553–561.

Yang G, Zhou X, Zhu J, Liu R, Zhang S, Coquinco A, Chen Y, Wen Y, Kojic L, Jia W, Cynader MS. 2013. JNK3 couples the neuronal stress response to inhibition of secretory trafficking. Sci Signal 6(283):ra57.

