## Supplementary material for "Palmitoylation Couples DLK to JNK3 to Facilitate Pro-degenerative Axon-to-Soma Signaling": JNK3 MS Suppl Figs n Legends

### Supplementary Figure Legends

**Fig S1. Schematic of different forms of JNK.** The products of the three JNK genes (JNK1-3; gene names *MAPK8-10*) are alternatively spliced, most notably to omit or include a non-catalytic C-terminal extension. The resultant forms migrate at approximately 46kDa (p46) and 54kDa (p54) on SDS-PAGE, respectively. Each form of JNK contains an identical Thr-Pro-Tyr motif that is dually phosphorylated by upstream MKKs to activate JNK (red circles with white 'P'). Modified from (Barr and Bogoyevitch 2001).

**Figure S2. Palmitoylation-dependence of DLK-JNK Signaling is Likely due to Direct Effects on DLK-JNK pathway kinases.** ***A:*** DRG neurons were treated with 2BrP (20 μM) or vehicle for 1h and then subjected to TD for 2.5h or left unstimulated in the continued presence of 2BrP or vehicle, as in Fig 2. Lysates were subjected to SDS-PAGE and subsequent immunoblotting to detect Akt phosphorylated at T308 (pAkt (T308)), total Akt, phosphorylated ERK (pERK) and total ERK. Right panel histogram confirms lack of effect of 2BrP on TD-induced dephosphorylation of Akt, n=3 determinations per condition (2-way ANOVA with Bonferroni *post hoc* test pAkt: 2BP p=0.90 [F(1)=0.017], TD p<0.0001 [F(1)=83.24], interaction p=0.84 [F(1)=0.041]. ***B:*** Lysates from *A* were immunoblotted to detect phosphorylated and total levels of ERK. Histograms confirm lack of effect of 2BrP on TD-induced dephosphorylation of ERK2-p42 or ERK1-p44 (middle and right histograms, respectively). ERK2-p42: 2BrP p=0.56 [F(1)=0.037], TD p<0.0001 [F(1)=62.66], interaction p=0.52 [F(1)=0.44]); ERK1-p44: 2BrP p=0.56 [F(1)=0.37], TD p<0.0001 [F(1)=62.66], interaction p=0.52 [F(1)=0.44]). ***C:*** Cultured DRG neurons were treated with 2BrP (20 μM) or EtOH vehicle and then incubated with Lysotracker dye. Representative kymographs reveal lysotracker-positive vesicles whose movement appears unaffected by 2BrP. Scale bar: 5μm. ***D:*** Quantified data from multiple time-lapse imaging experiments as in *C* confirm that 2BrP does not affect the proportion of lysotracker-positive vesicles that move anterogradely versus retrogradely (left panel histograms), does not affect the average velocity of retrograde or anterograde lysotracker-positive vesicles (center histograms) and does not affect the total number of lysotracker-positive vesicles per unit length of axon (right histograms). All data are shown as mean ± SEM.

**Figure S3. DLK mobility shift is due to phosphorylation.** GFP immunoprecipitates from lysates of HEK 293T cells cotransfected to express wtDLK-GFP and myc-JNK3 were treated with or without lambda phosphatase and subjected to SDS-PAGE. Phosphatase treatment increases DLK mobility.

**Figure S4. JNK1, JNK2 and JNK3 antibodies specifically recognize their intended targets.** Myc-tagged JNK1, JNK2 and JNK3 were expressed in HEK293T cells and anti-myc immunoprecipitates were blotted with the indicated antibodies. Anti-JNK1, -JNK2 and -JNK3 antibodies are all specific for their cognate antigens.

**Figure S5 Kinase dead mutation does not affect vesicular trafficking of wild type (palmitoylation-competent) JNK3.** Time-lapse images taken at the indicated times (top three panels) and kymograph (bottom panel) of axon from a DRG neuron expressing GFP-tagged kinase-dead JNK3 (GFP-JNK3 KD). Kinase-dead mutation (K55R) does not affect JNK3 axonal trafficking. Scale bar: 5 $\mu$m.

**Figure S6 Equal expression of AAV-delivered JNK3-WT and –CCSS in cultured DRG neurons.** Equivalent volumes of AAVs subsequently used in *in vivo* experiments in Figure 6 were first used to infect cultured DRG neurons. Immunoblotting of lysates harvested 5 days post-infection with anti-HA and anti tubulin (load control) antibodies confirms similar levels of total expression of HA-JNK3wt and HA-JNK3-CCSS.

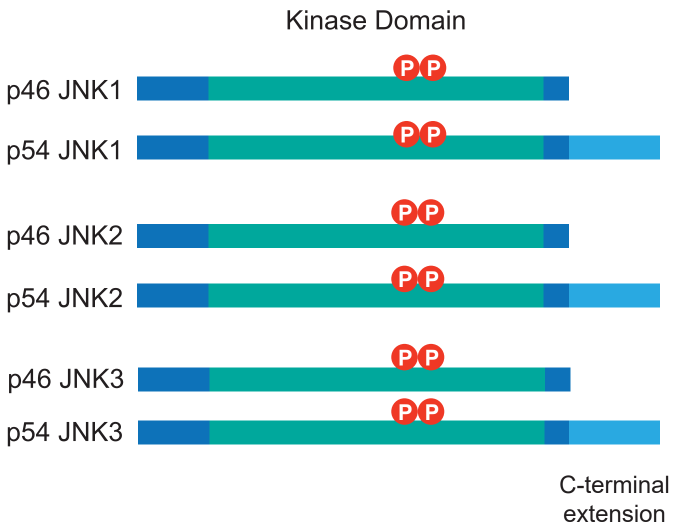


Niu et al., Figure S1

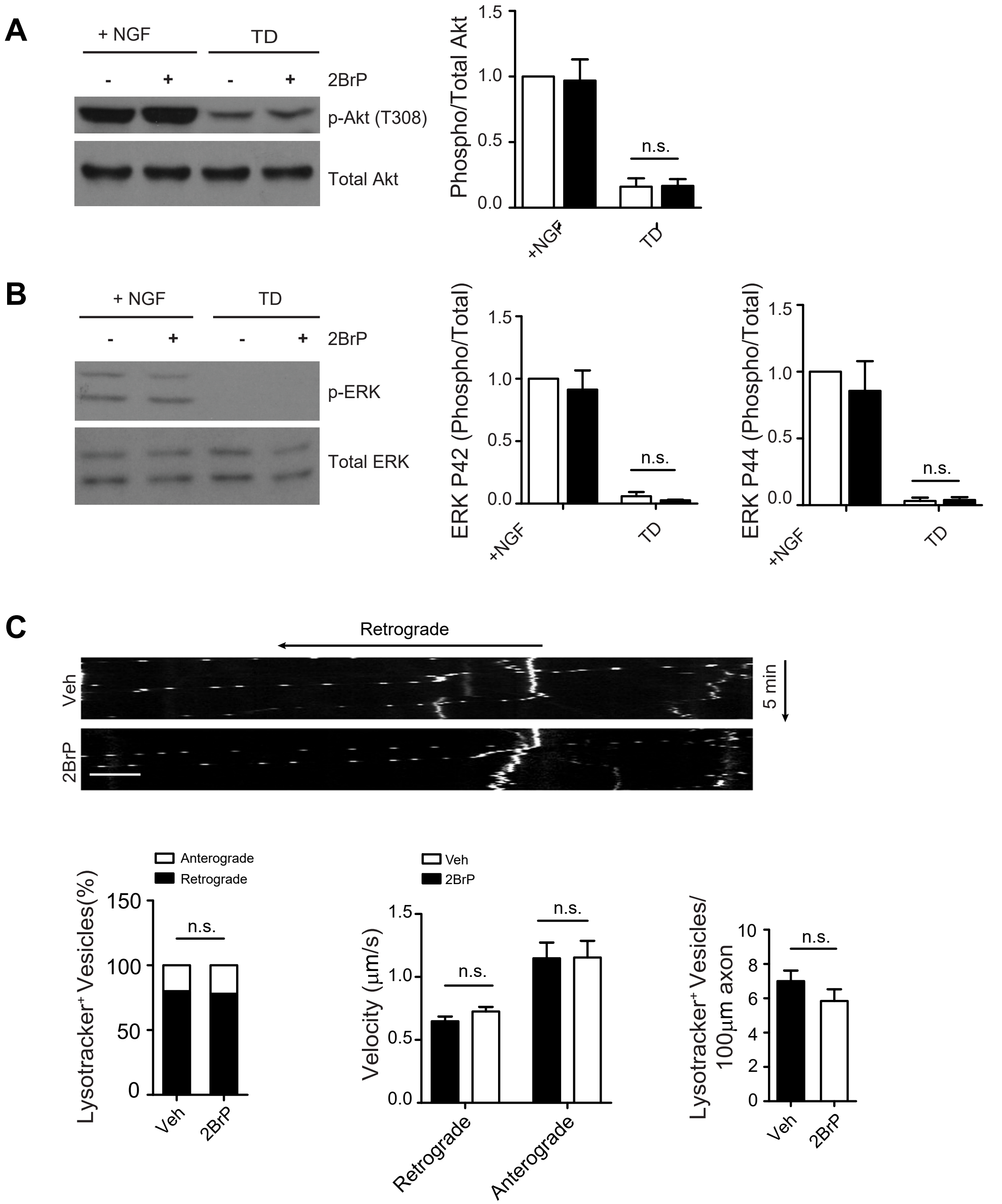


Niu et al., Figure S2

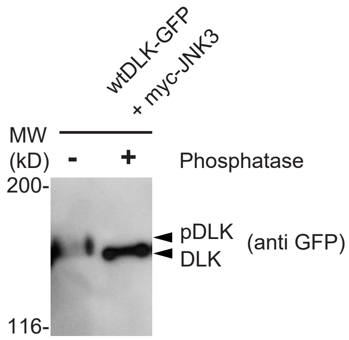
wtDLK-GFP
+ myc-JNK3
MW
(kD)
-
+
Phosphatase
200-
pDLK
DLK
(anti GFP)
116-

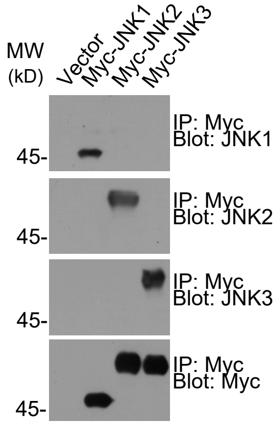


Niu et al., Figure S4

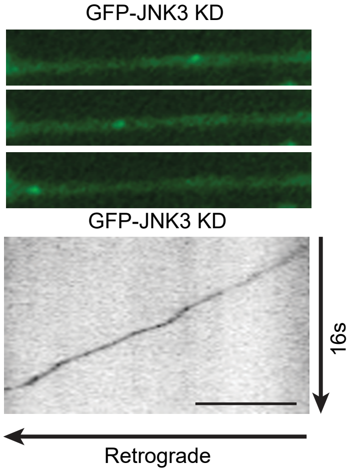


Niu et al., Figure S5

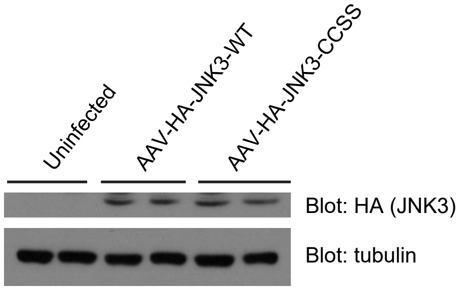
Uninfected
AAV-HA-JNK3-WT
AAV-HA-JNK3-CCSS
Blot: HA (JNK3)
Blot: tubulin
